# ChiCMaxima: a robust and simple pipeline for detection and visualization of chromatin looping in Capture Hi-C

**DOI:** 10.1101/445023

**Authors:** Yousra Ben Zouari, Anne M Molitor, Natalia Sikorska, Vera Pancaldi, Tom Sexton

## Abstract

Capture Hi-C (CHi-C) is a new technique for assessing genome organization, based on chromosome conformation capture coupled to oligonucleotide capture of regions of interest such as gene promoters. Chromatin loop detection is challenging, since existing Hi-C/4C-like analyses, which make different assumptions about the technical biases presented, are often unsuitable. We describe a new approach, ChiCMaxima, which uses local maxima combined with a background model to detect DNA looping interactions, integrating information from biological replicates. ChiCMaxima shows more stringency and robustness compared to previously developed tools. The tool includes a GUI browser for flexible visualization of CHi-C profiles alongside epigenomic tracks.

## Background

The advent of the chromosome conformation capture (3C) technology [1] allowed higher-order chromosome folding to be inferred by identifying spatial proximity between distal genomic sequences, leading to a comprehensive insight of genome topology. As sequencing throughput has increased, it has become feasible to globally assess all chromatin interactions within a population (4C: “one-to-all”; 5C: “many-to-many”; Hi-C: “all-to-all” methods) simply by sequencing all 3C ligation products or a selected subset of them [2-5]. In fact, Hi-C interaction maps can give insight into chromosome folding at different scales, depending on the sequencing depth (and hence resolution) of the study [6, 7]. However, the strength of Hi-C in assessing all possible chromatin interactions is also one of its major disadvantages: the numbers of possible ligation products that can be detected is much greater than current sequencing output. Recently, several groups have coupled Hi-C (or another 3C derivative) to sequence capture with pools of oligonucleotides complementary to thousands of restriction fragment ends [8-12]. Such “CHi-C” (Capture Hi-C) methods allow the simultaneous and higher resolution mapping of chromatin interactions for large subsets of the genome, such as all promoters or DNase hypersensitive sites. For example, promoter-centered interactomes have already been used to assign epigenomic status and follow enhancer looping dynamics throughout development, as well as to characterize disease-linked intergenic sequence polymorphisms [13-17]. Despite being highly informative, CHi-C datasets have specific properties that set them apart from other 3C-like techniques, which require specialized analytical tools to take these aspects into account. The majority of CHi-C strategies involve large numbers (thousands) of genomically dispersed baits for which interacting regions are detected. The asymmetry between the number of baits and the number of detected interacting regions leads to an asymmetry of CHi-C contact matrices, confounding standard Hi-C normalization approaches. In addition, individual baits have variable capture efficiencies which introduce additional technical biases. Depending on the bait design, CHi-C datasets will be more or less populated with ligation products between two bait fragments (“double-captured” interactions), as well as between bait and non-bait (“single-captured”), which may complicate bias assessment even further.

As for all genome-wide datasets, the challenges for CHi-C analysis are in the appropriate definition of an expected background level, from which “significant” signal can be resolved, and in the development of correct normalization strategies to reduce the impact of non-biological biases. Up to now, two major methods have been described for CHi-C analysis: GOTHiC [18] and CHiCAGO [19]. GOTHiC, actually developed for interaction calling in Hi-C, employs a very simplistic binomial test coupled with multiple testing correction to search for over-represented interactions, but does not explicitly take into account known features of Hi-C data, such as the heavy dependence of “background” interactions on genomic distance, let alone aspects of CHi-C such as capture bias. CHiCAGO uses a statistical background model to account for different biases in promoter-CHi-C data, combining three factors to define the expected background interaction level: genomic distance, bait capture efficiency, and technical biases present in Hi-C and sequencing approaches [19]. These parameters are fitted to the data to define an expected interaction strength for each individual restriction fragment, based on a combined negative binomial and Poisson variable. However, the treatment of each single fragment as an independent variable creates problems when accounting for biological replicates, since despite its improved coverage compared to Hi-C, current depths of CHi-C datasets still vastly sub-sample the possible space of ligation products. As a result, many reproducible chromatin loops observed at the resolution of larger bins of pooled restriction fragments are lost when scoring individual restriction fragments (Additional File 1: Figure S1). Related to this, it also follows that chromatin interactions comprising contiguous fragments of increased signal, centered on an interaction peak, are less likely to result from technical artefacts than isolated “spikes” of CHi-C signal. CHiCAGO utilizes the same geometric mean approach as DESeq2 [20] to allow weighting for different read depths of different replicates, but this may not completely counter the problem, especially if there is a large discrepancy in numbers of sequence reads between replicates. We tried to overcome these existing limitations of CHi-C analysis methods, and developed ChiCMaxima, which we applied to published mouse embryonic stem (ES) cell promoter CHi-C data [11]. Benchmarking against GOTHiC and CHiCAGO showed that ChiCMaxima was a more stringent method for interaction calling, but more robust to handling undersampling when comparing biological replicates. Further, ChiCMaxima gave a higher enrichment for interactions containing hallmarks of regulatory chromatin, such as histone modifications indicative of enhancers or CTCF binding sites, suggesting that its false positive detection rate for functional chromatin loops may be lower than for the other methods. Analysis of the chromatin contact network resulting from ChiCMaxima-called interactions identified potential key roles of Polycomb proteins and elongating RNA polymerase II, in line with previous findings [21], further demonstrating the utility of ChiCMaxima. In addition to the pipeline for calling CHi-C interactions, we also present ChiCBrowser, a user-friendly and flexible browser for inputting whole CHi-C datasets and then normalizing and visualizing bait-specific interaction profiles. Tracks of annotated genes and linear epigenomic profiles can also be added to the browser, and called interactions (whether by ChiCMaxima or other methods) can also be highlighted. This tool, whether used standalone or in parallel with ChiCMaxima interaction calling, will aid the community to analyze CHi-C datasets and inform new hypotheses.

## Results

### Methodological foundation of ChiCMaxima

#### Calling interactions as signal local maxima

In 3C approaches, genomic distance has an important impact on the expected frequency of interactions. Generally, the frequency of interactions decays with a power law as the genomic distance between fragments increases, consistent with many polymer physics models [4]. DNA loops correspond to a peak of higher interaction signal compared to the expected level of neighbor fragments on either side; this principle was used to detect loops in some of the first 3C studies [22]. To detect peaks in the signal, we use a naïve, non-parametric approach to call local maxima, without making any prior assumptions or using any preconceived model of the data (Fig. 1). The theoretical basis and proof of principle of ChiCMaxima is presented below; an operational guide and breakdown of the pipeline’s different tools is detailed in Additional File 2.

**Figure 1.**
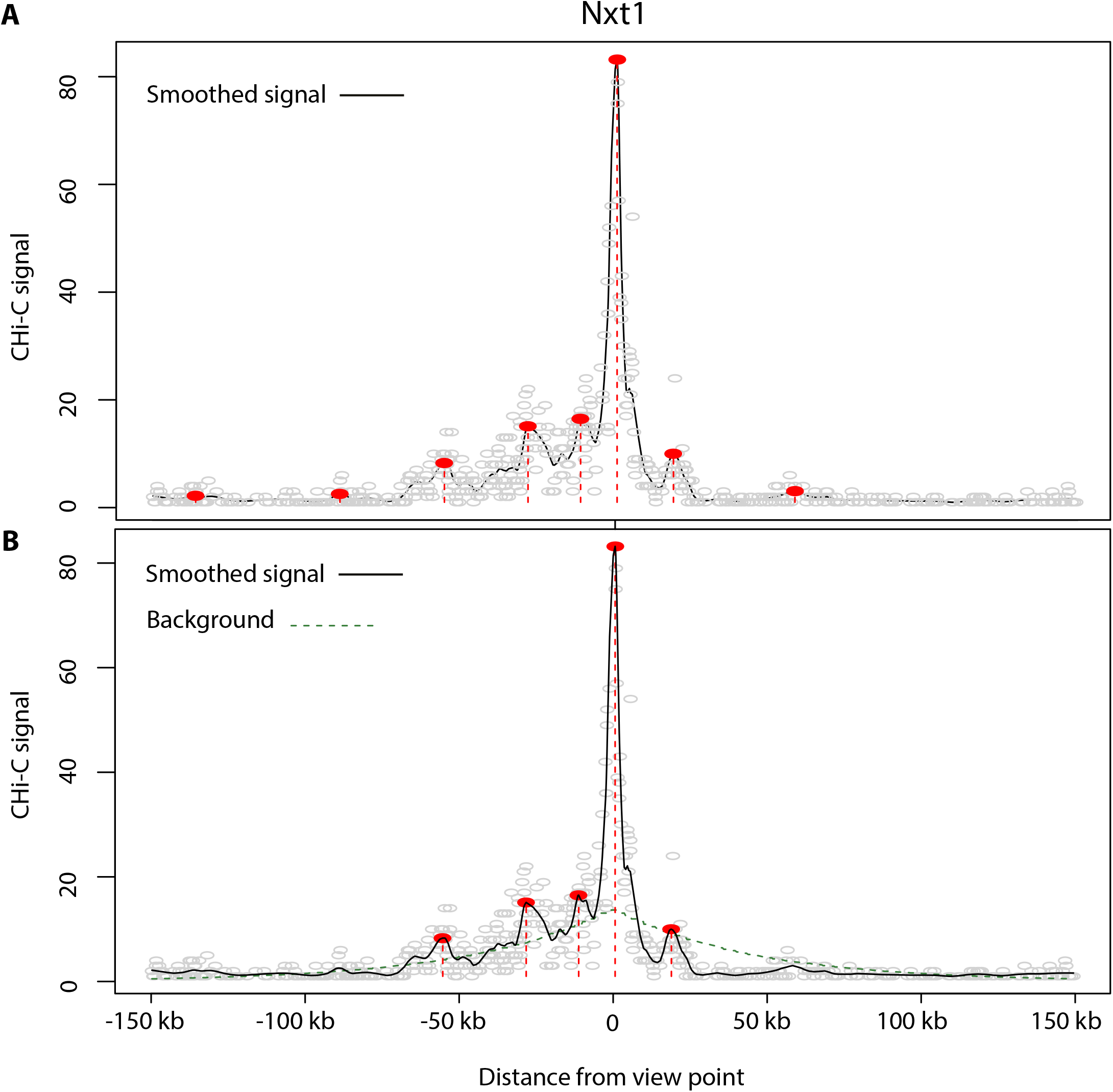
Interaction calling by ChiCMaxima. **a** Virtual 4C profile derived from one mES CHi-C replicate centered on the bait *Nxt1* promoter. The numbers of raw CHi-C sequence reads are plotted as gray circles against their genomic location, and the black line shows the loess-smoothed profile (span = 0.05). Red dotted lines and filled circles denote the positions of called interactions, defined as local maxima of smoothed signal within a fixed number of covered restriction fragments (window = 50). **b** The same virtual 4C profile as **a**, but with a dotted line (dark green) representing the estimated background signal as determined by a negative binomial fit to the bait-specific signal decay profile. Red dotted lines and filled circles denote the same local maxima as **a** which have smoothed signal higher than the estimated background.

First, treating each bait independently and removing bait-to-bait and inter-chromosomal interactions, we obtain a “virtual 4C” profile of read counts relative to the genomic position of the non-bait fragment, and perform loess smoothing on this profile. The fragments with the maximum signal are identified within sliding windows of a fixed fragment number, and local maxima are defined as regions where the smoothed signal equals this value. With this approach, only two parameters need to be controlled: the span of the loess smoothing (*s*), and the window size (*w*) for the local maximum computation. Over-smoothing or using too large a window size causes some maxima to be missed, and under-smoothing or small window sizes calls many local spikes as spurious interactions. ROC (receiver operating characteristic) analysis found smaller *s* and larger *w* to give more power in calling interactions, although greater numbers of bait-proximal restriction fragments need to be covered by sequence reads for local maximum computation as *w* increases (Additional File 1: Figure S2). CHi-C analysis in this manuscript is performed with the parameters *s* = 0.05, *w* = 50, a choice which optimizes interaction calling power and appropriate coverage of baits. Regardless of the choice of *s* and *w* parameters, we observed that local maxima with very low signal, which are very distant from the bait (and thus have a negligible background signal from neighboring fragments), are still called as “interactions” (Fig. 1a). We opted to filter out these spurious calls with an estimation of the background model.

#### Estimation of the background level

According to previous work on CHi-C data [19], the background interaction level at short genomic distances (up to ~1.5 Mb) is largely dominated by genomic separation (proposed to be caused by Brownian collisions of the chromosome fiber). In CHiCAGO, a cubic-fitted distance function was derived from the geometric means of read counts for binned genomic separations, and was then scaled with capture bias estimates in the final derived background distribution [19]. Inspired by this, we derived similar but *bait-specific* genomic distance functions, which are fitted to each virtual 4C profile. The advantage of this approach is that the data from different baits, which may reside in very different chromatin environments, do not need to be pooled together. The major limitation is that the relative paucity of bait-specific data could lead to overfitting in the model, particularly if a strong interaction causes overestimation of “background” signal around it. Instead of a cubic fit, we applied a fit to a negative binomial distribution, to account for the known overdispersion of sequencing data, and to provide a very conservative estimate of background [20]. Filtering ChiCMaxima calls to only those that exceeded this conservative background estimation was successful in removing the spurious, low-signal, distal local maxima that likely represent false positives (Fig. 1b). The ChiCMaxima tool outputs the log2 ratio of the local maximum signal versus the estimated background, which could be used as a means of ranking called interaction “strengths”. However, the possibility of model overfitting means that this approach should be used with caution; the major utility of background estimation is to remove spurious, low-signal local maxima.

#### Accounting for biological replicates

Although CHi-C improves on the resolution afforded by conventional Hi-C, it remains an under-sampled method. Although taking the intersection of called interactions from all replicates will give the highest-confidence chromatin loops, the false negative rate appears to be very high from this approach, due to poor reproducibility at the single restriction fragment level, both for CHiCAGO and for the better-performing ChiCMaxima (Additional File 1: Figure S1). We noted that many interaction peaks from one biological replicate also had adjacent or very close peaks in the second replicate, even though they were not at exactly the same restriction fragment (Additional File 1: Figure S3a). To see if these are likely to represent the same biological interactions, we assessed more systematically the distributions of genomic distance between interacting regions called in one biological replicate and the closest interaction called in the second replicate (Additional File 1: Figure S3b). Indeed, around one quarter of ChiCMaxima-called interactions had no genomic separation across replicates, meaning that they were on the same or directly contiguous restriction fragment, and half of all interactions were found within 20 kb (~5 *Hind*III restriction fragments), suggesting that genomic interactions called by CHi-C can indeed be reproducibly called across replicates, albeit at a lower resolution than single restriction fragments. To add more flexibility for analyzing biological replicates, ChiCMaxima allows a threshold distance between reported peaks in biological replicates to be defined by the user (*d*: default in the tool is 0). After local maximum computation and background model filtering on each biological replicate, these interactions are further filtered to retain only those where an interaction is also called within distance *d* in all other biological replicates. Unless stated otherwise, CHi-C analysis in this manuscript is performed with the parameter *d* = 20 kb. ChiCMaxima also provides a tool for assessing the distributions of closest distances between interactions called in pairs of biological replicates, better informing the user on their choice of the *d* parameter (see Additional File 2 for details).

#### Benchmarking of ChiCMaxima

We performed ChiCMaxima on a published mouse ES promoter CHi-C dataset [11], and compared our results with published ones from GOTHiC and CHiCAGO applied to the same dataset [11, 19] (Table 1; Additional File 3: Table S1). On visual inspection, ChiCMaxima successfully identified clear promoter interactions, some of which we also validated by 4C, and seemed to call fewer spurious ones than the other two methods (Fig. 2). Indeed, ChiCMaxima identified fewer promoter-centered interactions (22,222) than CHiCAGO (94,148) or GOTHiC (548,551). Pairwise comparisons revealed a striking dissimilarity of called interactions across all three methods - with the exception of ChiCMaxima interactions within the GOTHiC set, the majority of called interactions from one method is not shared with those of another (Fig. 3a). This is likely due to the very different assumptions made in the models for each method. We next sought to compare the performance of each method in calling chromatin interactions that are most likely to be functionally relevant, and minimizing likely false positives. First, we tested the hypothesis that ChiCMaxima, in calling fewer interactions than the other two methods, was the most stringent tool, calling only higher-confidence interactions. We split the interaction sets called by CHiCAGO or GOTHiC into those that were recapitulated, or not, by ChiCMaxima. In both cases, the interactions maintained in ChiCMaxima had significantly higher metrics of interaction score (weighted probability score in CHiCAGO [19]; observed/expected ratio in GOTHiC [18]) than for interactions called by the other method alone (Fig. 3b; *P* < 2×10^−16^, Wilcoxon rank sum test). Interactions conserved by CHiCAGO and GOTHiC calling also had significantly higher observed/expected ratios than interactions called in GOTHiC alone, but with a much more modest effect size. We thus conclude that ChiCMaxima is indeed the most stringent of the CHi-C interaction calling methods, calling the higher confidence interactions of the other methods.

**Table 1:** Overview of CHi-C interactions called by CHiCAGO, GOTHiC and CHiCMaxima.

|  | **CHiCAGO**<br>[19] | **GOTHic**<br>[11] | **ChiCMaxima**<br>(this<br>manuscript) | **ChiCMaxima**<br>and<br>**CHiCAGO** |
| --- | --- | --- | --- | --- |
| **Number of called interactions** | 94148 | 548551 | 22222 | 4089 |
| **Mean number of called interactions per bait** | 4.2 | 29.4 | 1.6 | 0.2 |
| **Median distance of intra-chromosomal interactions** | 155200 bp | 34776 bp | 133900 bp | 14390 bp |

**Figure 2.**
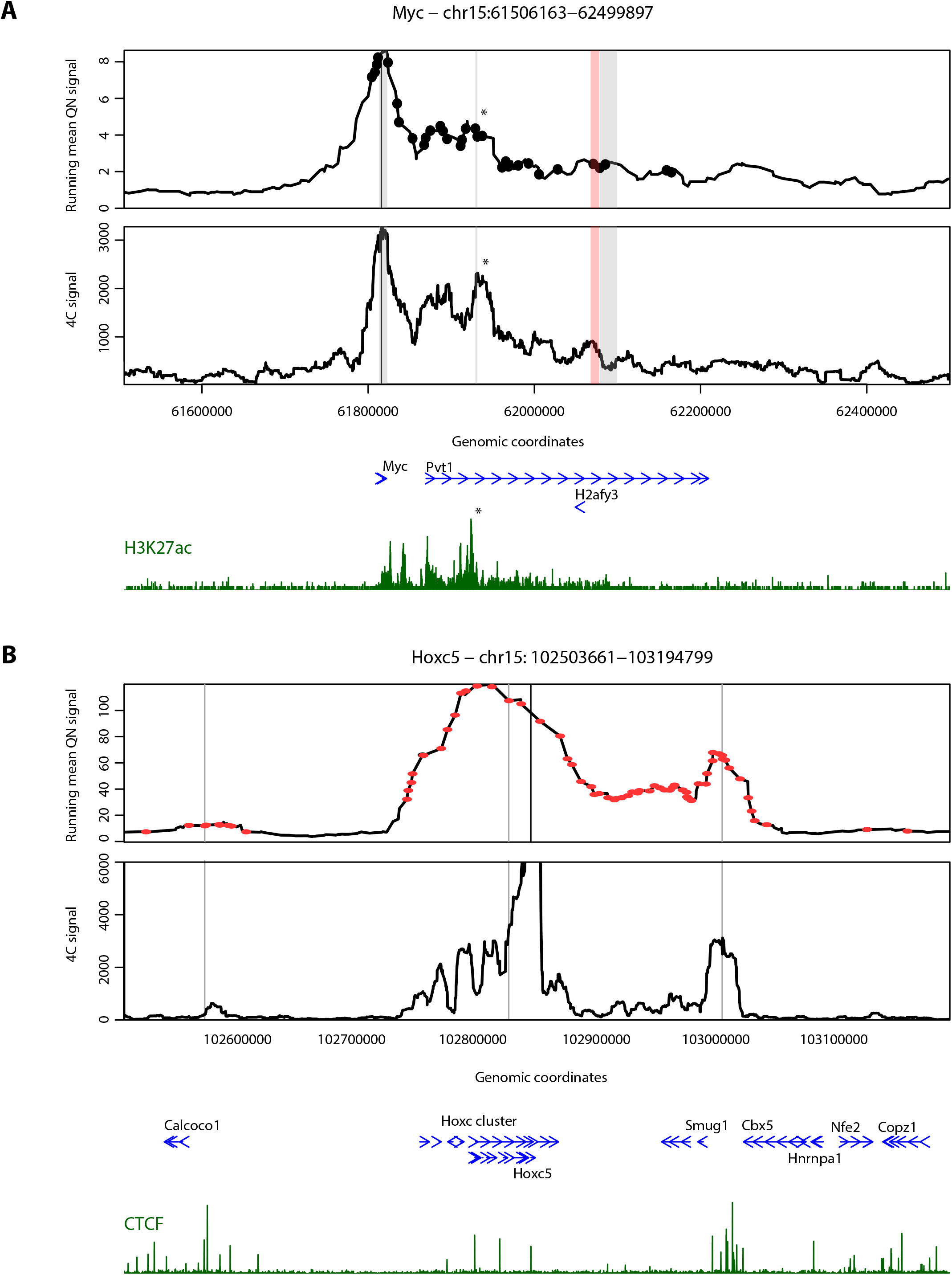
ChiCMaxima precisely calls chromatin interactions. **a** mES CHi-C (upper panel) and 4C (lower panel) profiles centered on the bait *Myc* promoter are shown. The interactions called by ChiCMaxima and CHiCAGO are denoted as stripes (gray and pink, respectively); points denote interactions called by GOTHiC. CHiCAGO missed an interaction between the *Myc* promoter and a putative enhancer marked by high H3K27ac (*); GOTHiC seemingly calls many spurious interactions. **b** mES CHi-C (upper panel) and 4C (lower panel) profiles centered on the bait *Hoxc5* promoter are shown. The interactions called by ChiCMaxima are denoted as gray stripes, and a large number of seemingly spurious interactions called by CHiCAGO are denoted as red points. Called interactions conserved between ChiCMaxima and CHiCAGO are centered on CTCF sites. For both profiles, gene position (blue) and ChIP-seq profiles (dark green; H3K27ac for **a**, CTCF for **b**) are shown below the CHi-C and 4C profiles.

**Figure 3.**
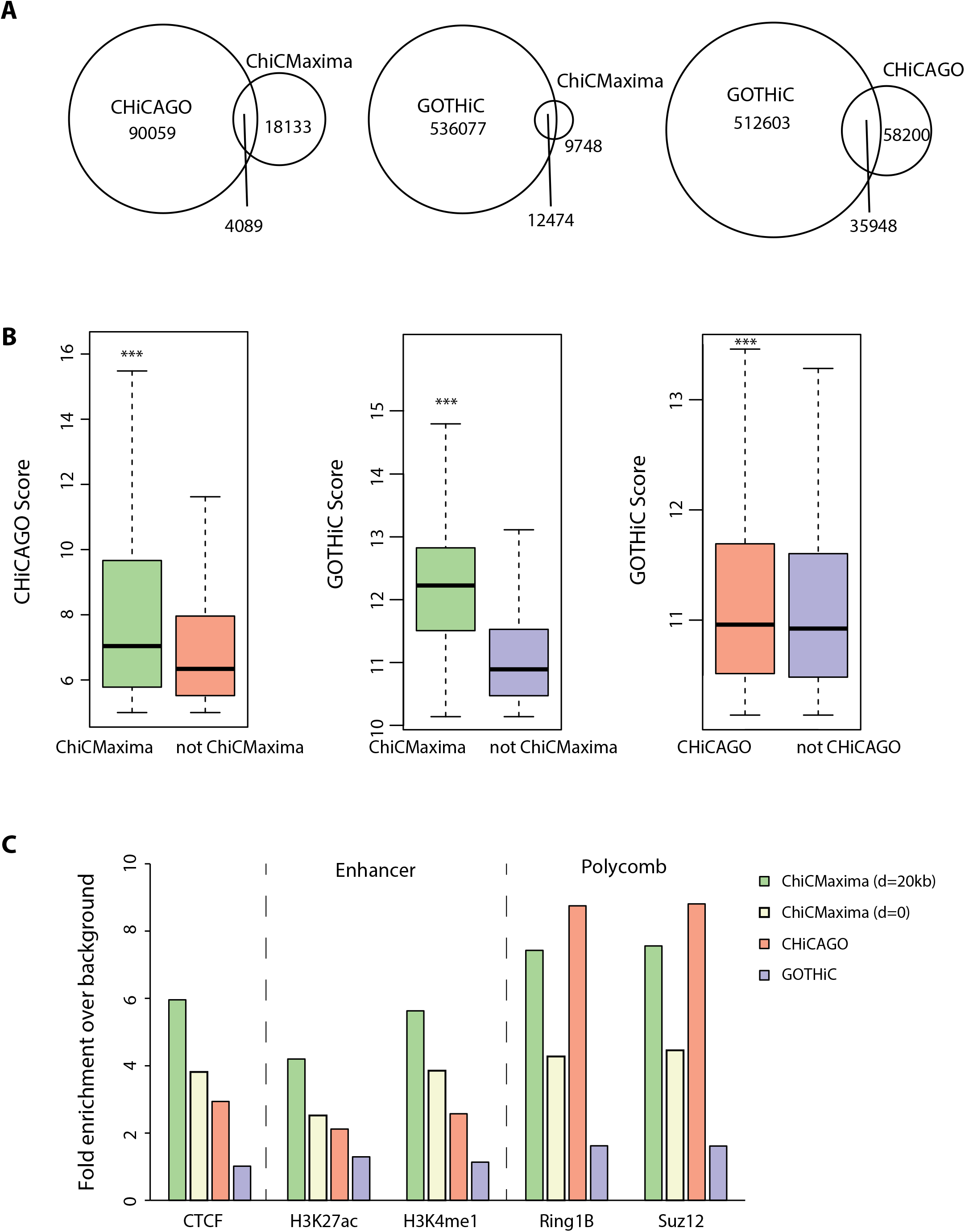
Comparison of ChiCMaxima, CHiCAGO and GOTHiC. **a** Venn diagrams showing numbers of interactions called by the three different methods which are conserved with the other methods. **b** Box plots comparing the CHiCAGO (left) or GOTHiC (center and right) metric scores of interaction strength for the sets of interactions called by CHiCAGO (left) or GOTHiC (center and right) which are conserved with those called by ChiCMaxima (left and center) or CHiCAGO (right), versus those which are not. *** *P* < 2×10^−16^; Wilcoxon rank sum test. **c** Bar charts showing fold enrichment over genomic background for different ChIP-seq peaks within the promoter-interacting sequences determined by the different CHi-C analysis methods.

#### Epigenomic analysis of ChiCMaxima-called interactions

One of the major perceived applications of CHi-C is to assign target genes to candidate *cis*- regulatory elements, particularly enhancers, by virtue of the specific interactions they make with promoters. Genomic studies revealed that enhancers share hallmark chromatin features: monomethylation of histone H3 lysine-4 (H3K4me1), DNase-hypersensitivity, acetylation of histone H3 lysine-27 (H3K27ac) and/or p300 co-activator occupancy [23]. However, despite epigenomic predictions of enhancers in numerous cell types, unambiguous identification of their target genes has proved more elusive, since they can control multiple genes, and may skip one or several promoters to act over large distances [24]. Promoter CHi-C studies have indeed shown a general enrichment in interacting regions bearing enhancer chromatin signatures [8, 10, 11], as well as for regions bound by CTCF, a known factor implicated in chromatin loops [25]. We reasoned that an interaction calling method that found the greatest proportion of putative enhancers and/or CTCF sites within a promoter CHi-C dataset was most likely to have the best true positive detection rate. Based on this, ChiCMaxima compares favorably to the other two methods. It has a higher enrichment for interacting regions containing CTCF, H3K27ac and H3K4me1 (Fig. 3c), with a ~2-fold improvement over CHiCAGO and ~5-fold improvement over GOTHiC. The enrichment in these functional hallmarks is decreased when the *d* parameter of ChiCMaxima is reduced to zero, but is still slightly better than CHiCAGO for enhancer marks. Chromatin interaction networks mediated by Polycomb group proteins have also been well described in embryonic stem cells [17, 21, 26, 27]. Reflecting this, promoter-interacting regions called by ChiCMaxima and CHiCAGO are also comparably and highly enriched in binding for core components of the two major Polycomb repressive complexes, Ring1B and Suz12 (Fig. 3c). Since more than half of CHiCMaxima-called interactions are not conserved with CHiCAGO, we also asked whether combining both methods would improve predictive power further. Indeed, the enrichment in functional hallmarks is even higher within the 4089 interactions that are conserved in both tools (Additional File 1: Figure S4a), indicating that combining the two methods gives the most stringent, highest-confidence interactions that are the most likely to be functionally relevant. However, the high enrichment for functional marks within ChiCMaxima-alone (and to a lesser extent for enhancer marks, CHiCAGO-alone) interactions implies that many functional interactions are also likely to be missed by intersecting the two methods. This is also apparent on visual inspection of called interactions within CHi-C profiles (Additional File 1: Figure S4b).

Furthermore, we assessed which of the 19,200 candidate mouse ES enhancers (based on chromatin signatures [28]) could be assigned to target promoters by the different methods (Table 2). As expected, the proportion of assigned enhancers scaled with the numbers of total called interactions (71.4% for GOTHiC; 19.2% for CHiCAGO; 14.5% for ChiCMaxima). However, candidate enhancers comprised a much higher proportion of the ChiCMaxima-called interaction set than for the other two methods (~3-fold higher than CHiCAGO; ~5-fold higher than GOTHiC), in line with the relative enrichments for individual regulatory marks. The interactions called by both ChiCMaxima and CHiCAGO only assign target genes to 3.4% of putative enhancers, with a modest increase in proportions of putative enhancers within the interaction set. Overall, these results suggest that ChiCMaxima provides a good compromise of stringency and coverage when assigning target genes to putative *cis*-regulatory elements.

**Table 2:**
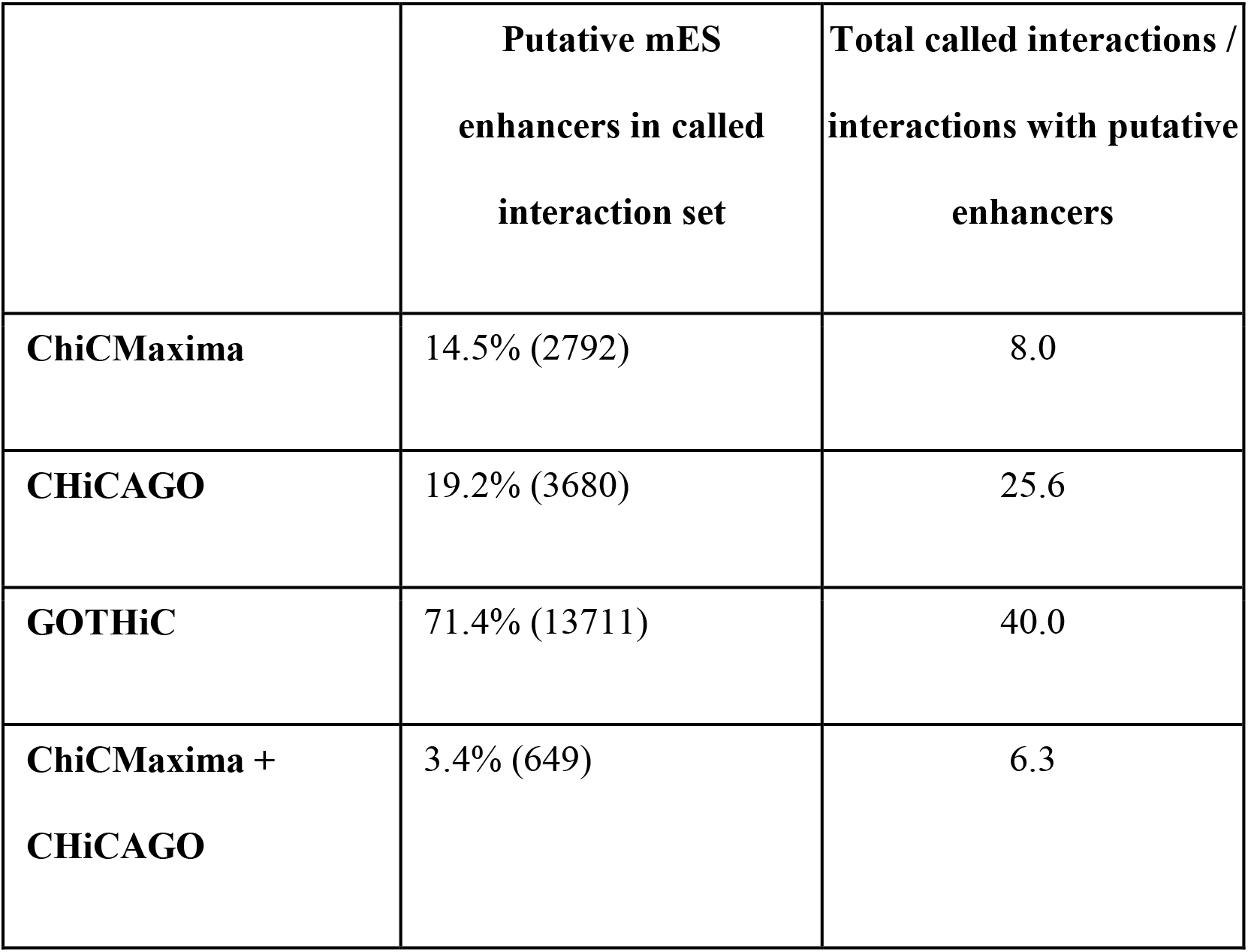
Overview of putative mES enhancers found within CHi-C interactions called by different methods.

|  | **Putative mES**<br>**enhancers in called**<br>**interaction set** | **Total called interactions /**<br>**interactions with putative**<br>**enhancers** |
| --- | --- | --- |
| **ChiCMaxima** | 14.5% (2792) | 8.0 |
| **CHiCAGO** | 19.2% (3680) | 25.6 |
| **GOTHiC** | 71.4% (13711) | 40.0 |
| **ChiCMaxima +**<br>**CHiCAGO** | 3.4% (649) | 6.3 |

### Chromatin assortativity analysis comparing CHiCAGO and ChiCMaxima derived contact networks

Despite great progress in the experimental mapping of chromatin organization inside the nucleus, many questions regarding the functional impact of its structure remain unanswered. It is thus difficult to estimate the accuracy of any interaction calling algorithm beyond the performance on the few regions of the genome that are well characterized. Moreover, alongside the known role of interactions in bringing enhancer regions close to their target genes and grouping Polycomb repressed genes, there might be other functionally relevant 3D chromatin structures which we still do not understand. Hence the need for finding complementary analytical methods to study the panorama of genome-wide interactions. A recent step in this direction was made by looking at chromatin contact maps as networks and applying methods from network theory to gain a comprehensive understanding of nuclear organization [e.g. 21, 29-31]. For example, appreciation of the chromatin interaction network topology bolstered the link between spatial gene co-associations and their co-expression patterns [29]. An important concept that was recently applied to chromatin interaction networks is *assortativity* – which indicates the extent to which genomic regions sharing the same chromatin mark(s) preferentially interact. This property is not trivially related to the relative abundance of a mark at interacting regions, and highly assortative chromatin features are more likely to be related to chromatin interactions. A recent study of mouse ES chromatin interactions identified three major chromatin features that were highly assortative: the abundant H3K4me1 mark, features of transcriptional elongation (predominantly RNA polymerase II phosphorylated on serine-2 of the C-terminal repeat domain (RNAPII-S2P) and trimethylation of lysine-36 of histone H3 (H3K36me3)), and the relatively low abundance Polycomb group proteins and associated histone marks (e.g. trimethylation of lysine-27 of histone H3 (H3K27me3) [21]. To further test the utility of ChiCMaxima, we applied chromatin assortativity (ChAs) analysis to the network of ChiCMaxima-called interactions, and directly compared it to the one derived by CHiCAGO for promoter-other end interactions (Fig. 4; Additional File 1: Figure S5). Although the relative abundances of the different chromatin features were very similar (Pearson correlation coefficient 0.99), and the three aforementioned categories of assortative chromatin features were identified by the two methods (Pearson correlation coefficient 0.89 for ChAs values obtained on the two networks), some differences were apparent. Whereas CHiCAGO identifies a slightly more robust Polycomb network, transcriptional elongation hallmarks are very strongly flagged by ChiCMaxima. In addition to RNAPII-S2P and H3K36me3, other features enriched within active gene bodies in ES cells, such as dimethylation of histone H3 lysine-79 (H3K79me2) and CBX3 (HP1γ; associated with transcriptional elongation and stem cell identity [32, 33]) were also revealed to be highly assortative by ChiCMaxima. These results demonstrate that ChiCMaxima-called interactions can be used in informative network analyses and highlight promoter-gene body contacts as a potentially important architectural feature for active genes (see Discussion).

**Figure 4.**
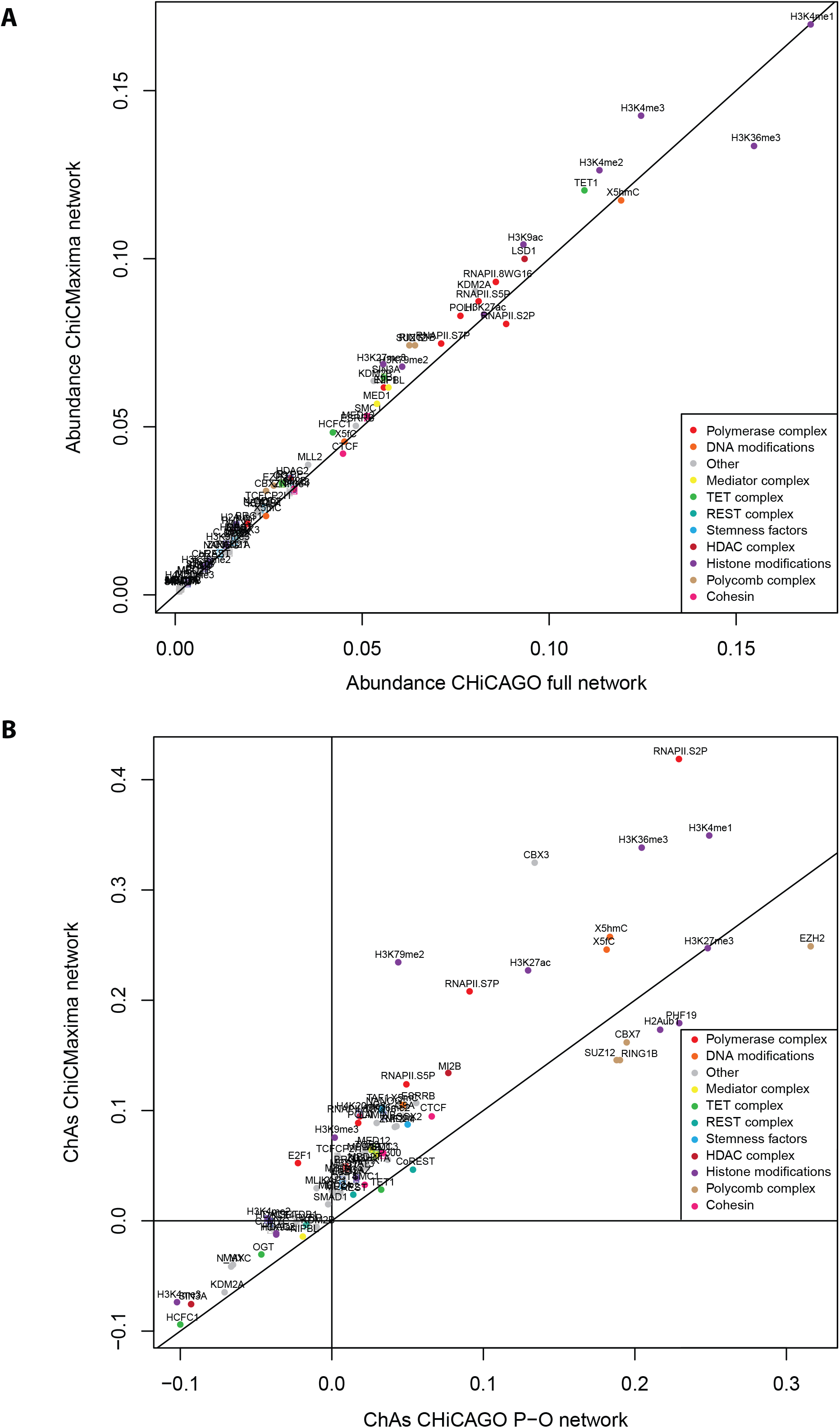
Exploration of chromatin assortativity of different features on the ChiCMaxima generated chromatin contact. **a** Scatter plot of abundance of different chromatin features within the interaction networks called by ChiCMaxima or CHiCAGO. **b** Scatter plot of chromatin assortativity of different chromatin features within the interaction networks called by ChiCMaxima or CHiCAGO (restricted to promoter-other end interactions). The class of the different chromatin features is color coded.

### ChiCBrowser

To enable visualization of promoter (and other sparse bait) CHi-C results, alongside linear epigenomic profiles and the interactions called by ChiCMaxima or other methods, we also developed ChiCBrowser, an R-based GUI browser. Unlike the WashU browser [34], which displays all interactions simultaneously and can be difficult to interpret visually, ChiCBrowser displays virtual 4C profiles, with the bait and display window defined by the user via a graphical window (Fig. 5). Its major functionalities are described below; a full user guide is given in Additional File 2.

**Figure 5.**
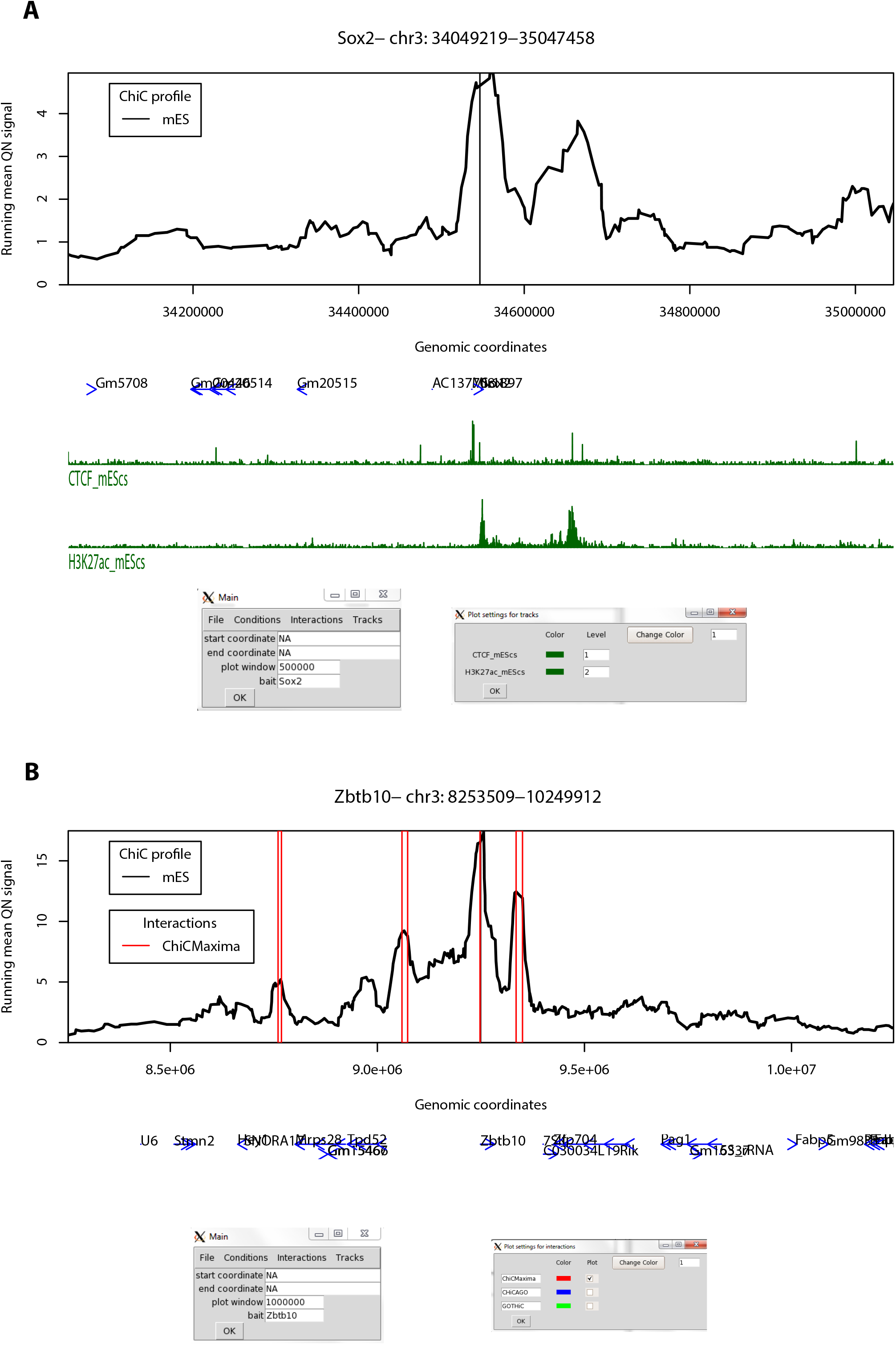
Some functionalities of ChiCBrowser. **a** A screenshot of ChiCBrowser, showing the mES CHi-C profile for 500 kb up- and downstream of the bait *Sox2* promoter. Gene positions (blue) and selected mES ChIP-seq tracks (green) are shown below the profile. The main ChiCBrowser user interface window is shown underneath (left), where the bait and plot window have been specified. A sub-window, called from the *Tracks* menu (right), allows the color and level of the epigenomic profiles to be controlled by the user. Epigenomic tracks that are given the same level (for instance, the same histone mark in different tissues) are scaled to the same level on the y-axis so that the profiles are visually comparable. **b** As for **a**, a screenshot of the mES CHi-C profile for 1 Mb up- and downstream of the bait *Zbtb10* promoter. Open red rectangles show the position of interactions called by ChiCMaxima. The sub-menu on the bottom right, called from the *Interactions* menu, allows the user to control which interaction lists to annotate on the CHi-C plot.

All CHi-C datasets which may be plotted together or compared are made into one input file (see Additional File 2 for input format details), which only needs to be loaded once into the memory for all subsequent plots to be made. To allow fairer comparisons between datasets, all CHi-C-derived virtual 4C profiles are quantile normalized [35] before the running mean values are plotted. Since CHi-C experimental designs usually include biological replicates and different conditions to be compared, ChiCBrowser provides flexibility (via the *Conditions* menu) to define the plot levels of each single CHi-C dataset. For example, (shown in Additional File 1: Fig. S6), biological replicates can be allocated to different levels and plotted side by side to compare experimental reproducibility, or given the same plot level, so that the mean profile can be plotted for comparison with other experimental conditions. The user can assign names to these plot levels and change their plotting colors.

The *Tracks* menu allows the user to load gene annotations (as a modified bed file; see Additional File 2), which are plotted as blue arrows to show transcriptional orientation, and linear epigenomic profiles in bigWig or bedGraph formats. Similarly to the *Conditions*, the user can define plot levels for epigenomic profiles (Fig. 5a). In this case, this defines which profiles are scaled to the same level on the y-axis, for instance allowing fairer comparison between profiles of the same histone mark mapped in different tissue types. The epigenomic profile plot colors can also be modified by the user.

Ostensibly, the *Interactions* menu allows the user to load sets of interactions called by ChiCMaxima (or CHiCAGO, whose output is in the same format) for them to be highlighted on the CHi-C profile (Fig. 5b). However, the input format of these interactions is essentially the chromosomal coordinates of genomic regions associated with a specific bait (see Additional File 2 for details), so this plotting functionality can be adapted to highlight any subset of the CHi-C dataset that the user designs (e.g. interactions unique to one condition or tissue type and not another). This flexibility in particular makes ChiCBrowser very useful to explore different hypotheses when browsing interactomes. As for other ChiCBrowser functions, the user can alter the name and color of these annotations, as well as select or de-select subsets of them.

## Discussion

We present two tools for processing and interpretation of Chi-C datasets: ChiCMaxima for interaction calling, and ChiCBrowser for bait-specific visualization of interaction profiles. Both were developed to overcome the currently identified unique challenges presented by these data. Despite a clear improvement over conventional Hi-C with limited sequencing throughput, the main issue with CHi-C outputs is that they are greatly under-sampled, creating problems of reproducibility across biological replicates at the highest resolutions (Additional File 1: Figures S1 and S3). The subsequent paucity of bait-specific data confounds the generation of powerful statistical models, so previous methods either appear to have high false positive rates (e.g. GOTHiC; see Fig. 2), and/or rely on combining data from multiple baits (e.g. CHiCAGO) to avoid overfitting model parameters. ChiCMaxima minimizes the number of model parameters to be estimated by naively just searching for local maxima in the virtual 4C profiles (Fig. 1a), a logic for calling chromatin loops that was used in some of the first 3C studies [22, 36]. Derivation of a background model, accounting for chromatin interaction decay with increasing genomic separation, was necessary to remove spurious local maxima in distal regions of low signal, and we opted for the conservative negative binomial model applied to bait-specific data. Despite concerns of overfitting, this approach performed well (Fig. 1b). For single datasets, only two parameters need to be defined in ChiCMaxima: the loess smoothing span (which consistently performs well when set to 0.05), and the window for local maximum computation. The power of peak calling is greater for larger windows, with the tradeoff that deeper sequence coverage is required (Additional File: Figure S2),. Thus a limited tuning of this parameter is still required from the user to find the optimum values for their own datasets. A major advantage of ChiCMaxima is thus that interactions can simply be called without the need to control or estimate multiple parameters, or choose arbitrary thresholds. However, this advantage also means that ChiCMaxima does not return measurements of statistical significance interpretable as the interaction “strength” of the called chromatin loops. The ratio of the local maximum signal to the expected background signal is returned, but this cannot be taken as a reliable measure of interaction strength (for example, not correlating with CHiCAGO interaction scores; data not shown). When comparing chromatin interactions between different tissues or conditions, we find that quantile normalization allows fair visual comparisons (e.g. Additional File: Figure S1 and S3), but further work will be required to better define and quantify interaction strength differences.

As mentioned previously, another major challenge resulting from the undersampling of CHi-C data is the handling of biological replicates. Presumably because it processes sliding windows rather than treating each restriction fragment independently, ChiCMaxima has superior reproducibility to CHiCAGO, but this is still less than 10% at the single restriction fragment level (Additional File: Figure S1). Since many interactions are reproduced at slightly lower resolutions (Additional File: Figure S3), ChiCMaxima has a built-in flexibility whereby interactions can be filtered for those that are conserved in all replicates, within a user-defined distance. The optimal distance is likely to vary between experiments, particularly with sequencing depth and complexity of the assessed genome. For this reason we provide tools to allow the user to explore the distributions of closest distances between interactions called in pairs of replicates and thus determine the optimal setting.

Despite the simplistic approach of ChiCMaxima, it compares favorably to GOTHiC and CHiCAGO in various different benchmarks, suggesting that it is one of the more stringent calling methods (thus likely reducing false positives) to successfully call a high proportion of interactions that are likely to be functionally relevant (Fig. 3). This includes tests of: reproducibility between biological replicates; increased metrics of interaction strength within ChiCMaxima-called interactions; enrichment for putative enhancer marks, CTCF binding sites and Polycomb-bound regions within promoter-interacting regions; assignment of putative enhancers to target genes; proportion of putative enhancers within the called interaction set; reduced apparent false positive rate on visual inspection of CHi-C profiles (e.g. Fig. 2). We note that promoter interactions with non-enhancer/CTCF/Polycomb-bound elements may certainly be frequent and functionally significant, albeit poorly characterized so far. Indeed, all three methods call many interactions of this category. However, the greater enrichment of ChiCMaxima-called interactions for promoter-enhancer loops that have been so well described in the literature, coupled with their overall higher interaction score metrics as called by other methods, suggests that ChiCMaxima is the most stringent interaction calling method, but also reliably identifies interactions most likely to be functionally relevant. ChiCMaxima also performs mostly favorably or equally as well as CHiCAGO when the most stringent setting for handling biological replicates is used (*d* = 0). However, the apparent inconsistency in interaction calls between the three methods (Fig. 3a), coupled with the good enrichment for regulatory marks in CHiCAGO-only interactions, suggests that ChiCMaxima has some false negatives which are correctly detected by CHiCAGO (the inverse also seems to be the case). Indeed, the highest-confidence interactions are conserved between CHiCAGO and ChiCMaxima, but the false negative rate seems very high when relying on this stringent approach. Overall, we recommend using ChiCMaxima when looking for global features of chromatin interactions, since the false positive rate seems lower, but combinations of ChiCMaxima and CHiCAGO may be required to comprehensively explore the interactomes of specific baits of interest. We also note that ChiCMaxima, due to its dependence on searching for local maxima, is not suitable for assessing ultra-long-range (> 2 Mb) or *trans* interactions, where the background signal is too low for local maxima to be reliable. Bait-to-bait interactions should also not be assessed by ChiCMaxima, since these double-captured interactions are highly likely to appear as “artificial” local maxima when flanked by single-captured, bait-to-non-bait interactions within sliding windows. Finally, CHi-C strategies using tiled oligonucleotides to intensively cover a contiguous domain [9, 37] are better analyzed with the suite of tools adapted to the contact matrices generated by 5C or Hi-C (e.g. adaptations of my5C [38] or Juicer [39]).

As a further demonstration of the utility of ChiCMaxima, network analysis of called chromatin interactions also identified the Polycomb-mediated interactome that has been previously described in ES cells [17, 21, 26, 27] (Fig 4). Interestingly, the ChiCMaxima network also indicates frequent contacts between promoters and the bodies of active genes, a phenomenon which was also identified by the same analysis of the CHiCAGO network, but to a lesser extent [21]. It is currently unclear whether this may be an indirect effect of transcriptional elongation on topology of the chromosome fiber [40], or reflects more specific mechanisms of gene expression control. For example, enhancers have been described to initially contact promoters, but to additionally track along the gene during transcriptional elongation [41], and promoter and enhancer interactions with specific exons have been implicated in splicing control [42, 43]. Further studies will be required to determine the functional significance, if any, of such intragenic chromatin looping events, but ChiCMaxima seems to be a very useful tool for studying them via CHi-C studies.

The ChiCBrowser tool is a flexible, user-friendly GUI to generate virtual 4C profiles, necessary for visual inspection of most CHi-C datasets. It has a built-in flexibility to allow biological replicates or different combinations of biological conditions to be assessed in parallel, and a similar flexibility is also built into the management of gene annotations and epigenomic profiles that are plotted alongside the CHi-C data (Fig. 5). Called interactions, whether by ChiCMaxima or other methods, can be easily highlighted on the display, based on a simple input format that can be adapted to highlight any subset of the CHi-C subset that may be of interest to the user. Overall, this browser will be of use to anyone wishing to explore CHi-C data.

## Conclusions

Capture Hi-C, particularly strategies with sparse baits such as promoters, is a rapidly growing technique hampered by the limited tools available to meet the unique challenges of analyzing the datasets produced. ChiCMaxima adopts a simplistic approach, with minimal prior assumptions on the data, and successfully calls CHi-C interactions, performing favorably with existing methods in various benchmarks. Most notably, ChiCMaxima provides the flexibility to deal with problems of reproducibility across biological replicates at high resolutions, a persistent but often overlooked challenge of CHi-C. Combined with the user-friendly, flexible ChiCBrowser, we provide a suite of tools for CHi-C analysis and visualization which will be of use to many in the nuclear organization community.

## Methods

### Datasets used in this study

Accession numbers and linked references for the CHi-C and ChIP-seq sequencing data, as well as pre-processed lists of interaction calls and putative mES enhancers, can be found in Additional File 1: Table S2.

### Sample pre-processing

Raw sequencing reads were processed by custom perl and R scripts, originally derived from the Hi-C analysis pipeline developed in [44], which entails mapping the paired reads with Bowtie [45], pairing, removing common Hi-C artefacts (PCR duplicates, circularized fragments, non-digested fragments) and then converting from genomic coordinates to restriction fragment space. Operationally, this generates only tiny differences from outputs of HiCUP [46]. Custom perl scripts, explained in Additional File 2 and available on https://github.com/yousra291987/ChiCMaxima, were used to convert paired bed files to the input format for ChiCMaxima. These scripts can also be applied to outputs of other Hi-C analysis tools, such as HiCUP [46], HiC-Pro [47] or Juicer pre-inputs [39].

### ChiCMaxima

The suite of scripts, made for R version >= 3.2, and its full documentation (including package dependencies, found on Bioconductor or CRAN), is available on https://github.com/yousra291987/ChiCMaxima. A full description of its usage, and how it is run on supplied test data, is also provided in Additional File 2. In brief, ChiCMaxima_Caller identifies interactions as local maxima of loess smoothed bait-specific interaction profiles within single CHi-C datasets. ChiCMaxima_RepAnalysis determines the distributions of the closest distance between interactions called in pairs of datasets, allowing the user to select an optimal threshold for filtering “maintained” interactions within biological replicates. ChiCMaxima_MergeRep2 or ChiCMaxima_MergeRepMany then applies this set distance threshold to identify interactions that are conserved in two or more biological replicates, respectively. Finally, ChiCMaxima_Collate is a utility script that generates one large table from multiple CHi-C datasets, convenient for input into ChiCBrowser. Except where stated specifically in the text, ChiCMaxima_Caller was run on each ES CHi-C replicate with the parameters *window_size* = 50, *loess_span* = 0.05, *cis_window* = 1500000; ChiCMaxima_MergeRep2 was run on their outputs with the parameter *repdist* = 20000.

### ChiCBrowser

The browser is run from an R environment (version >= 3.2), and its full documentation (including package dependencies, found on Bioconductor or CRAN), is also available on https://github.com/yousra291987/ChiCMaxima. A full user guide is also presented in Additional File 2, along with examples of its use on supplied test data. This browser, or small variants in the code (e.g. to show raw data instead of after smoothing by running means in Additional File: Figure S1b) were used to generate all the screenshot images presented in the article.

### CHiCAGO and GOTHiC interaction lists

The previously called lists of interactions from both mES CHi-C replicates using CHiCAGO (GSE81503_mESC_PCHiC_merge_final_washU_text.txt; CHiCAGO score >= 5) or GOTHiC (ESC_promoter_other_significant_interactions.txt; log (observed/expected) >= 10) were downloaded directly from their repositories (see Additional File 1: Table S2). For interaction calling within individual biological replicates by CHiCAGO, interactions with a score >= 5 were used after running CHiCAGO with default parameters (maxLBrownEst = 1500000; minFragLen = 150; maxFragLen = 40000; minNPerBait = 250; binsize = 20000; removeAdjacent = TRUE; adjBait2bait = TRUE; tlb.filterTopPercent = 0.01; tlb.minProxOEPerBin = 50000; tlb.minProxB2BPerBin = 2500; techNoise.minBaitsPerBin = 1000; brownianNoise.samples = 5; brownianNoise.subset = 1000; brownianNoise.seed = NA; weightAlpha = 34.11573; weightBeta = -2.586881; weightGamma = -17.13478; weightDelta = -7.076092). Intersections of called interactions across biological replicates (Additional File 1: Figure S1a) were found by searches for called interactions with identical Bait_name and ID_OE columns.

### ROC analysis and assessing ChiCMaxima w parameter limits

For different values of *w* and *s*, a variant of ChiCMaxima_Caller, which made no corrections for estimations of background (see Fig 1a), was run on one replicate on a random subset of 1000 baits, comprising a total dataset of > 200,000 bait-to-non-bait combinations with covered sequence reads. For all of the assessed bait-to-non-bait combinations, a table was compiled of the number of supporting sequence reads (N) and a binary calling score (1 if the interaction was called by this instance of ChiCMaxima_Caller, 0 if not). These two variables were then input into the R package ROCR [48], with the performance assessed for “true positive rate” versus “true negative rate”, with the “average” threshold set. For assessment of the limits of the *w* parameter (Additional File 1: Figure S2b), the number of non-bait fragments covered by sequencing reads within 1.5 Mb of the interacting bait were counted for each bait; for ChiCMaxima_Caller to assess local maxima with a bait, this number must be equal to or greater than 2*w* + 1.

### Assessing distances between potentially conserved interactions across biological replicates

This is performed by ChiCMaxima_RepAnalysis. Pairs of interaction files (the output of ChiCMaxima_Caller) are split according to their bait, and one set is defined as the *query* and the other set as the *subject*. For each non-bait fragment within the query, the genomic distance to the closest non-bait fragment within the subject set is found by the utilities within the R GenomicRanges package [49].

### 4C interaction validation

J1 mouse ES cells were grown on gamma-irradiated mouse embryonic fibroblast cells under standard conditions (4.5 g/L glucose-DMEN, 15% FCS, 0.1 mM non-essential amino acids, 0.1 mM beta-mercaptoethanol, 1 mM glutamine, 500 U/mL LIF, gentamicin), then passaged onto feeder-free 0.2% gelatin-coated plates for at least two passages to remove feeder cells. Cells were detached with trypsin, washed by centrifugation in PBS, and then fixed with 2% formaldehyde in mES culture medium for 10 min at 23°C. The fixation was quenched with cold glycine at a final concentration of 125 mM, then cells were washed with PBS and permeabilized on ice for 1 h with 10 mM Tris-HCl, pH 8, 100 mM NaCl, 0.1% NP-40 and protease inhibitors. Nuclei were resuspended in *Dpn*II restriction buffer at 10 million nuclei/mL concentration, and 5 million nuclei aliquots were further permeabilized by treatment for 1 h with 0.4% SDS at 37°C, then a further 1h with 2.6% Triton-X100 at 37°C. Nuclei were digested overnight with 1000 U *Dpn*II at 37°C, then washed twice by centrifuging and resuspending in T4 DNA ligase buffer. *In situ* ligation was performed in 400 μL T4 DNA ligase buffer with 20,000 U T4 DNA ligase overnight at 16°C. DNA was purified by reverse cross-linking with an overnight incubation at 65°C with proteinase K, followed by RNase A digestion, phenol/chloroform extraction and isopropanol precipitation. The DNA was digested with 5 U/μg *Csp*6I at 37°C overnight (for *Myc*) or 5 U/μg *Tai*I at 65°C for 2 h (for *Hoxc5*), then re-purified by phenol/chloroform extraction and isopropanol precipitation. The DNA was then circularized by ligation with 200 U/μg T4 DNA ligase under dilute conditions (5 ng/μL DNA), and purified by phenol/chloroform extraction and isopropanol precipitation. 50 ng aliquots of this DNA were used as template for PCR with bait-specific primers containing Illumina adapter termini (primer sequences and optimal PCR conditions available on request). PCR reactions were pooled, primers removed by washing with 1.8x AMPure XP beads, then quantified on a Bioanalyzer (Agilent) before sequencing with a HiSeq 4000 (Illumina). Sequence reads were filtered and mapped to *Dpn*II restriction fragments, essentially as previously described [5, 50]. For visualization of the 4C profiles, running means of read counts across windows of 25 restriction fragments are plotted against the genomic coordinate of the fragment interacting with the bait (Fig 2).

### Comparing CHi-C calling methods

The intersections in interaction calling methods (Fig 3a) were computed using the R GenomicRanges package [49] to find overlapping coordinates within the non-bait regions from interaction sets with the same bait. Comparisons of the interaction scores from CHiCAGO or GOTHiC-called interactions which were or were not conserved with another method were computed by Wilcoxon rank sum tests.

### Assessing enrichment for epigenomic marks

ChIP-seq fastq files (see Additional File 1: Table S2) were aligned to the mm9 genome with bowtie2 [45], then peaks were called with the Erange 4.0 ChIP-seq peak finder tools [51, 52], with the settings --nodirectionality, --notrim and an FDR threshold of 0.05. Enrichment of each epigenetic feature within an interaction set was computed by dividing the proportion of interactions (non-bait component) overlapping with a feature peak within the interaction set by the proportion of all mappable, non-bait restriction fragments which overlap with a feature peak. These overlaps were found using bedtools [53] on the bed files of non-bait interacting regions versus the bed files of called ChIP-seq peaks. Overlaps of the set of putative mES enhancers [28] with non-bait regions within called interactions were performed with GenomicRanges.

### Chromatin assortativity

Interaction network analysis was performed exactly as described in [21]. Briefly, 78 chromatin features were taken from [54] and peak-calling/binarization was performed as described there in 200 bp windows. For each fragment the overlapping windows of chromatin peaks were identified and their values averaged to give a fraction of presence of any feature in each fragment. The abundance of a feature is defined as the average of that feature value across all fragments in the network considered. ChAs of a specific chromatin feature is defined as the Pearson correlation coefficient of the value of that feature across all pairs of nodes that are connected with each other. They are computed from the “assortativity” function of the R package igraph. ChAs were computed from the total interaction network derived by ChiCMaxima, which omits bait-to-bait interactions. To avoid confounding effects of bait-to-bait interactions present within the full CHiCAGO-called network, ChAs computation was restricted to only the promoter-to-other end (P-O) portion of the network.

## Declarations

### Ethics approval and consent to participate

Not applicable.

### Consent for publication

Not applicable.

### Availability of data and materials

The ChiCMaxima tools and ChiCBrowser, along with supporting documentation and test data, are available on https://github.com/yousra291987/ChiCMaxima. 4C datasets have been submitted to GEO with the accession number (*will be submitted before publication*).

### Authors’ contributions

YBZ and TS conceived and developed the ChiCMaxima and ChiCBrowser tools and performed the benchmarking with other methods. AMM and NS performed and analyzed 4C experiments. VP performed assortativity and other chromatin interaction network analysis. All authors contributed to writing the manuscript.

## Acknowledgements

We thank all other members of the Sexton lab for advice. Sequencing was performed by the IGBMC GenomEast platform, a member of the France Genomique consortium (ANR-10- INBS-0009).

## References

1. Dekker J, Rippe K, Dekker M, Kleckner N. Capturing chromosome conformation. Science. 2002;2955558:1306–11.

2. Dostie J, Richmond TA, Arnaout RA, Selzer RR, Lee WL, Honan TA, et al. Chromosome Conformation Capture Carbon Copy (5C): a massively parallel solution for mapping interactions between genomic elements. Genome research. 2006;1610:1299–309.

3. Fullwood MJ, Liu MH, Pan YF, Liu J, Xu H, Mohamed YB, et al. An oestrogen-receptor-alpha-bound human chromatin interactome. Nature. 2009;4627269:58–64.

4. Lieberman-Aiden E, van Berkum NL, Williams L, Imakaev M, Ragoczy T, Telling A, et al. Comprehensive mapping of long-range interactions reveals folding principles of the human genome. Science. 2009;3265950:289–93.

5. van de Werken HJ, Landan G, Holwerda SJ, Hoichman M, Klous P, Chachik R, et al. Robust 4C-seq data analysis to screen for regulatory DNA interactions. Nature methods. 2012;910:969–72.

6. Bonev B, Mendelson Cohen N, Szabo Q, Fritsch L, Papadopoulos GL, Lubling Y, et al. Multiscale 3D Genome Rewiring during Mouse Neural Development. Cell. 2017;1713:557–72 e24.

7. Rao SS, Huntley MH, Durand NC, Stamenova EK, Bochkov ID, Robinson JT, et al. A 3D Map of the Human Genome at Kilobase Resolution Reveals Principles of Chromatin Looping. Cell. 2014;1597:1665–80.

8. Hughes JR, Roberts N, McGowan S, Hay D, Giannoulatou E, Lynch M, et al. Analysis of hundreds of cis-regulatory landscapes at high resolution in a single, high-throughput experiment. Nature genetics. 2014;462:205–12.

9. Kolovos P, van de Werken HJ, Kepper N, Zuin J, Brouwer RW, Kockx CE, et al. Targeted Chromatin Capture (T2C): a novel high resolution high throughput method to detect genomic interactions and regulatory elements. Epigenetics & chromatin. 2014;7:10.

10. Sahlen P, Abdullayev I, Ramskold D, Matskova L, Rilakovic N, Lotstedt B, et al. Genome-wide mapping of promoter-anchored interactions with close to single-enhancer resolution. Genome biology. 2015;16:156.

11. Schoenfelder S, Furlan-Magaril M, Mifsud B, Tavares-Cadete F, Sugar R, Javierre BM, et al. The pluripotent regulatory circuitry connecting promoters to their long-range interacting elements. Genome research. 2015;254:582–97.

12. Andrey G, Schopflin R, Jerkovic I, Heinrich V, Ibrahim DM, Paliou C, et al. Characterization of hundreds of regulatory landscapes in developing limbs reveals two regimes of chromatin folding. Genome research. 2017;272:223–33.

13. Javierre BM, Burren OS, Wilder SP, Kreuzhuber R, Hill SM, Sewitz S, et al. Lineage-Specific Genome Architecture Links Enhancers and Non-coding Disease Variants to Target Gene Promoters. Cell. 2016;1675:1369–84 e19.

14. Mifsud B, Tavares-Cadete F, Young AN, Sugar R, Schoenfelder S, Ferreira L, et al. Mapping long-range promoter contacts in human cells with high-resolution capture Hi-C. Nature genetics. 2015;476:598–606.

15. Rubin AJ, Barajas BC, Furlan-Magaril M, Lopez-Pajares V, Mumbach MR, Howard I, et al. Lineage-specific dynamic and pre-established enhancer-promoter contacts cooperate in terminal differentiation. Nature genetics. 2017.

16. Siersbaek R, Madsen JGS, Javierre BM, Nielsen R, Bagge EK, Cairns J, et al. Dynamic Rewiring of Promoter-Anchored Chromatin Loops during Adipocyte Differentiation. Molecular cell. 2017;663:420–35 e5.

17. Joshi O, Wang SY, Kuznetsova T, Atlasi Y, Peng T, Fabre PJ, et al. Dynamic Reorganization of Extremely Long-Range Promoter-Promoter Interactions between Two States of Pluripotency. Cell stem cell. 2015;176:748–57.

18. Mifsud B, Martincorena I, Darbo E, Sugar R, Schoenfelder S, Fraser P, et al. GOTHiC, a probabilistic model to resolve complex biases and to identify real interactions in Hi-C data. PloS one. 2017;124:e0174744.

19. Cairns J, Freire-Pritchett P, Wingett SW, Varnai C, Dimond A, Plagnol V, et al. CHiCAGO: robust detection of DNA looping interactions in Capture Hi-C data. Genome biology. 2016;171:127.

20. Love MI, Huber W, Anders S. Moderated estimation of fold change and dispersion for RNA-seq data with DESeq2. Genome biology. 2014;1512:550.

21. Pancaldi V, Carrillo-de-Santa-Pau E, Javierre BM, Juan D, Fraser P, Spivakov M, et al. Integrating epigenomic data and 3D genomic structure with a new measure of chromatin assortativity. Genome biology. 2016;171:152.

22. Palstra RJ, Tolhuis B, Splinter E, Nijmeijer R, Grosveld F, de Laat W. The beta-globin nuclear compartment in development and erythroid differentiation. Nature genetics. 2003;352:190–4.

23. Heintzman ND, Hon GC, Hawkins RD, Kheradpour P, Stark A, Harp LF, et al. Histone modifications at human enhancers reflect global cell-type-specific gene expression. Nature. 2009;4597243:108–12.

24. Sanyal A, Lajoie BR, Jain G, Dekker J. The long-range interaction landscape of gene promoters. Nature. 2012;4897414:109–13.

25. Phillips JE, Corces VG. CTCF: master weaver of the genome. Cell. 2009;1377:1194–211.

26. Denholtz M, Bonora G, Chronis C, Splinter E, de Laat W, Ernst J, et al. Long-range chromatin contacts in embryonic stem cells reveal a role for pluripotency factors and polycomb proteins in genome organization. Cell stem cell. 2013;135:602–16.

27. Schoenfelder S, Sugar R, Dimond A, Javierre BM, Armstrong H, Mifsud B, et al. Polycomb repressive complex PRC1 spatially constrains the mouse embryonic stem cell genome. Nature genetics. 2015;4710:1179–86.

28. Chen CY, Morris Q, Mitchell JA. Enhancer identification in mouse embryonic stem cells using integrative modeling of chromatin and genomic features. BMC genomics. 2012;13:152.

29. Babaei S, Mahfouz A, Hulsman M, Lelieveldt BP, de Ridder J, Reinders M. Hi-C Chromatin Interaction Networks Predict Co-expression in the Mouse Cortex. PLoS computational biology. 2015;115:e1004221.

30. Sandhu KS, Li G, Poh HM, Quek YL, Sia YY, Peh SQ, et al. Large-scale functional organization of long-range chromatin interaction networks. Cell reports. 2012;25:1207–19.

31. Norton HK, Emerson DJ, Huang H, Kim J, Titus KR, Gu S, et al. Detecting hierarchical genome folding with network modularity. Nature methods. 2018;152:119–22.

32. Sridharan R, Gonzales-Cope M, Chronis C, Bonora G, McKee R, Huang C, et al. Proteomic and genomic approaches reveal critical functions of H3K9 methylation and heterochromatin protein-1gamma in reprogramming to pluripotency. Nature cell biology. 2013;157:872–82.

33. Vakoc CR, Mandat SA, Olenchock BA, Blobel GA. Histone H3 lysine 9 methylation and HP1gamma are associated with transcription elongation through mammalian chromatin. Molecular cell. 2005;193:381–91.

34. Zhou X, Lowdon RF, Li D, Lawson HA, Madden PA, Costello JF, et al. Exploring long-range genome interactions using the WashU Epigenome Browser. Nature methods. 2013;105:375–6.

35. Ritchie ME, Phipson B, Wu D, Hu Y, Law CW, Shi W, et al. limma powers differential expression analyses for RNA-sequencing and microarray studies. Nucleic acids research. 2015;437:e47.

36. Tolhuis B, Palstra RJ, Splinter E, Grosveld F, de Laat W. Looping and interaction between hypersensitive sites in the active beta-globin locus. Molecular cell. 2002;106:1453–65.

37. Franke M, Ibrahim DM, Andrey G, Schwarzer W, Heinrich V, Schopflin R, et al. Formation of new chromatin domains determines pathogenicity of genomic duplications. Nature. 2016;5387624:265–9.

38. Lajoie BR, van Berkum NL, Sanyal A, Dekker J. My5C: web tools for chromosome conformation capture studies. Nature methods. 2009;610:690–1.

39. Durand NC, Shamim MS, Machol I, Rao SS, Huntley MH, Lander ES, et al. Juicer Provides a One-Click System for Analyzing Loop-Resolution Hi-C Experiments. Cell systems. 2016;31:95–8.

40. Lavelle C. Pack, unpack, bend, twist, pull, push: the physical side of gene expression. Current opinion in genetics & development. 2014;25:74–84.

41. Lee K, Hsiung CC, Huang P, Raj A, Blobel GA. Dynamic enhancer-gene body contacts during transcription elongation. Genes & development. 2015;2919:1992–7.

42. Mercer TR, Edwards SL, Clark MB, Neph SJ, Wang H, Stergachis AB, et al. DNase I-hypersensitive exons colocalize with promoters and distal regulatory elements. Nature genetics. 2013;458:852–9.

43. Ruiz-Velasco M, Kumar M, Lai MC, Bhat P, Solis-Pinson AB, Reyes A, et al. CTCF-Mediated Chromatin Loops between Promoter and Gene Body Regulate Alternative Splicing across Individuals. Cell systems. 2017;56:628–37 e6.

44. Sexton T, Yaffe E, Kenigsberg E, Bantignies F, Leblanc B, Hoichman M, et al. Three-dimensional folding and functional organization principles of the Drosophila genome. Cell. 2012;1483:458–72.

45. Langmead B, Trapnell C, Pop M, Salzberg SL. Ultrafast and memory-efficient alignment of short DNA sequences to the human genome. Genome biology. 2009;103:R25.

46. Wingett S, Ewels P, Furlan-Magaril M, Nagano T, Schoenfelder S, Fraser P, et al. HiCUP: pipeline for mapping and processing Hi-C data. F1000Research. 2015;4:1310.

47. Servant N, Varoquaux N, Lajoie BR, Viara E, Chen CJ, Vert JP, et al. HiC-Pro: an optimized and flexible pipeline for Hi-C data processing. Genome biology. 2015;16:259.

48. Sing T, Sander O, Beerenwinkel N, Lengauer T. ROCR: visualizing classifier performance in R. Bioinformatics. 2005;2120:3940–1.

49. Lawrence M, Huber W, Pages H, Aboyoun P, Carlson M, Gentleman R, et al. Software for computing and annotating genomic ranges. PLoS computational biology. 2013;98:e1003118.

50. de Wit E, Vos ES, Holwerda SJ, Valdes-Quezada C, Verstegen MJ, Teunissen H, et al. CTCF Binding Polarity Determines Chromatin Looping. Molecular cell. 2015;604:676–84.

51. Johnson DS, Mortazavi A, Myers RM, Wold B. Genome-wide mapping of in vivo protein-DNA interactions. Science. 2007;3165830:1497–502.

52. Mortazavi A, Williams BA, McCue K, Schaeffer L, Wold B. Mapping and quantifying mammalian transcriptomes by RNA-Seq. Nature methods. 2008;57:621–8.

53. Quinlan AR, Hall IM. BEDTools: a flexible suite of utilities for comparing genomic features. Bioinformatics. 2010;266:841–2.

54. Juan D, Perner J, Carrillo de Santa Pau E, Marsili S, Ochoa D, Chung HR, et al. Epigenomic Co-localization and Co-evolution Reveal a Key Role for 5hmC as a Communication Hub in the Chromatin Network of ESCs. Cell reports. 2016;145:1246–57.

